# Optoacoustic imaging of GLP-1 Receptor with a near-infrared exendin-4 analog

**DOI:** 10.1101/2020.04.29.068619

**Authors:** Sheryl Roberts, Eshita Khera, Crystal Choi, Tejas Navaratna, Jan Grimm, Greg M. Thurber, Thomas Reiner

## Abstract

Limitations in current imaging tools have long challenged the imaging of small pancreatic islets in animal models. Here, we report the first development and in vivo validation testing of a broad spectrum and high absorbance near infrared optoacoustic contrast agent, E4_x12_-Cy7. Our near infrared tracer (E4_x12_-Cy7) is based on the amino acid sequence of exendin-4 and targets the glucagon-like peptide-1 receptor (GLP-1R). Cell assays confirmed that E4_x12_-Cy7 has a high binding affinity (IC_50_ = 4.6 ± 0.8 nM). Using the multi-spectral optoacoustic tomography (MSOT), we imaged E4_x12_-Cy7 and optoacoustically visualized ß-cell insulinoma xenografts *in vivo* for the first time. In the future, similar optoacoustic tracers that are specific for ß-cells and combines optoacoustic and fluorescence imaging modalities could prove to be important tools for monitoring the pancreas for the progression of diabetes.

## Introduction

Imaging does not presently play a prominent role in diabetes diagnostics. The current diagnosis for type 1 or 2 diabetes and prediabetes is based on the patient’s medical history and blood screening in the form of an AC1 test^1^. However, the prevalence of diabetes is expected to drive the demand and use of molecular imaging in order to facilitate the diagnosis and monitoring of the disease. Type 1/2 diabetes for all age-groups worldwide is projected to rise from 171 million in 2000 to 366 million in 2030^2^. The rise in the incident of diabetes-a major cause of blindness, kidney failure, heart attacks, stroke and lower limb amputation^3^-has not been met with a comparable, parallel refinement in diagnostic tools for its early detection. Unlike computed tomography (CT), magnetic resonance imaging (MRI), ultrasonography and intravital microscopy^4, 5^, which have played a major roles, in the field of diabetes imaging, optoacoustic imaging is thus far underrepresented among diagnostic tools. Optoacoustic glucose sensing techniques in blood and plasma samples, much like the AC1 test, have been reported as early as in 1999^6^. Few studies, however, focus on newly developed techniques of optoacoustic spectrometry and specific spectral unmixing algorithms intended to sense glucose^7–9^. While glucose sensing remains the most effective method for the detection of diabetes^10^, the direct visualization of ß-cells remains a high priority, as it promises to diagnose occult diabetes before metabolic imbalance and dysregulated blood levels manifest.

Βeta-cells are key regulators of glucose *balance via* secretion of insulin, and ß-cell dysfunction leads to chronic diseases, type 1 and type 2 diabetes^11, 12^. Prolonged high blood sugar levels can damage major organs, moreover and lead to the development of diseases and complications beyond diabetes^13, 14^. Although, molecular imaging of ß-cell function has always garnered interest, accurate *in vivo* ß-cell mass (BCM) identification and quantification *via* molecular imaging remains elusive for three main reasons: (1) the tracer of choice must be specific for ß-cells. (2) ß-cells make up only 1-2% of the pancreas, and (3) ß-cells are located deep in the pancreas, which itself is located between major organs that are involved in clearance pathways (liver, spleen and nested in between parts of the GI tract), precluding easy access for imaging. Furthermore, intravital microscopy requires invasive surgery due to light scattering, and positron emission tomography (PET) requires the use of radioactive materials. Thus far, *in vivo* intravital imaging and PET imaging methods have played only preclinical^15–17^ and early-phase clinical roles in studying the dynamic processes of pancreatic tissue^18, 19^.

Among other well-studied receptors^20–22^, glucagon-like peptide-1 receptors (GLP-1R) is abundant on the ß-cell surface and controls for blood sugar levels through insulin secretion^11, 12^. Since its discovery, GLP-1R has been a constant target for developing drugs. One such drug, exendin-4 (exenatide), well-known for its high affinity and efficacy is an excellent agonist ^4, 23^. Over the past few years, several fluorescent imaging agents based on exendin-4 have been described in the literature ^24–27^.

Alternatively, then, non-invasive optoacoustic tomography, which combines light and sound, could represent a viable option able to image at greater depths than other optical techniques. BCM need a reliable method for non-invasive in vivo imaging and quantification^28, 29^. A handful of suitable and clinically translatable two-dimensional (2D) and three-dimensional (3D) hand-held optoacoustic devices exists^30, 31^, including commercial options for clinical research^32–34^. Based on previous approaches and motivated by the undeniable clinical need to non-invasively resolve ß-cells mass *in vivo*, we explored the use of exendin as a targeted vector using multi-spectral optoacoustic tomography (MSOT), a preclinical optoacoustic platform, similar to the equipment used in the clinics. Ideally, acoustic signals from the agent would overcome endogenous background signal against the inherently small populations of ß-cells. We have shown in previous studies that a number of near-infrared (NIR) dyes are suitable for optoacoustic imaging ^35, 36^. Such NIR scaffolds are characteristically high absorbers (typically ε > 200, 000 cm^−1^ M^−1^) with a low quantum yield (θ) of highly packed *π*-conjugated aromatic systems and overlapping p-orbitals of delocalized electrons. Here, we explore the scope of NIR-exendin for applications in optoacoustic diabetes detection.

We aimed to synthesize a NIR optoacoustic sonophore, with site-specific labeling at a single position capable of targeting the GLP-1R. To this end, we conjugated an exendin-4 analog to Cy7 *via* copper-catalyzed click chemistry, replacing the K_12_ position of the amino acid sequence with the azide-reactive (S)-2-amino-4-pentynoic acid ^4, 26^, to synthesize E4_x12_-Cy7. In order to characterize E4_x12_-Cy7, we used tissue mimicking phantoms and mouse insulinoma xenografts that mimic normal expressions and function of GLP-1R akin to pancreatic ß-cells. The research described within this article incorporates our synthetic design to target GLP-1R on the surface of ß-cells using MSOT, representing the first optoacoustic visualization of insulinoma *in vivo.*

## Results

### Synthesis and analysis of E4_x12_-Cy7

A GLP-1R targeted peptide, exendin-4, was modified at the K_12_ position of the amino acid sequence with the unnatural alkyne-amino acid (S)-2-amino-4-pentynoic acid (Figure 1A). As reported previously^4, 24, 26^, it can be used without significant reduction of peptide/GLP-1R target binding (~3 nM)^4^. Likewise, we have previously shown that cyanine dyes such as Cy7 exhibit the desired optical characteristics suitable for both optoacoustic and fluorescent imaging^36^. The synthesis of E4_x12_-Cy7 was achieved in a one-step reaction (Figure 1B). Cy7-azide (Lumiprobe) was used for labeling of the neopeptide E4_x12_ (grey) via a bioorthogonal copper(I)-catalyzed azide-alkyne cycloaddition reaction (CuAAC) to obtain E4_x12_-Cy7 (Figure 1B, *bottom*) in good yields (>90%) and purity (78%, t_R_ = 12.15 min, Figure 1C). The targeting peptide exendin-4, neopeptide E4_x12_, and optoacoustic agent E4_x12_-Cy7 are amino acid sequences consisting of 39 amino acids (Figure 1A). E4_x12_-Cy7 has a broad absorption, ranging from 550-850 nm, with a maximum absorption at 760 nm, indicating an appropriate wavelength range for multi-spectral optoacoustic unmixing^37^. Mass spectrometry confirmed the identity of E4_x12_-Cy7 (Figure 1D).

**Figure 1.**
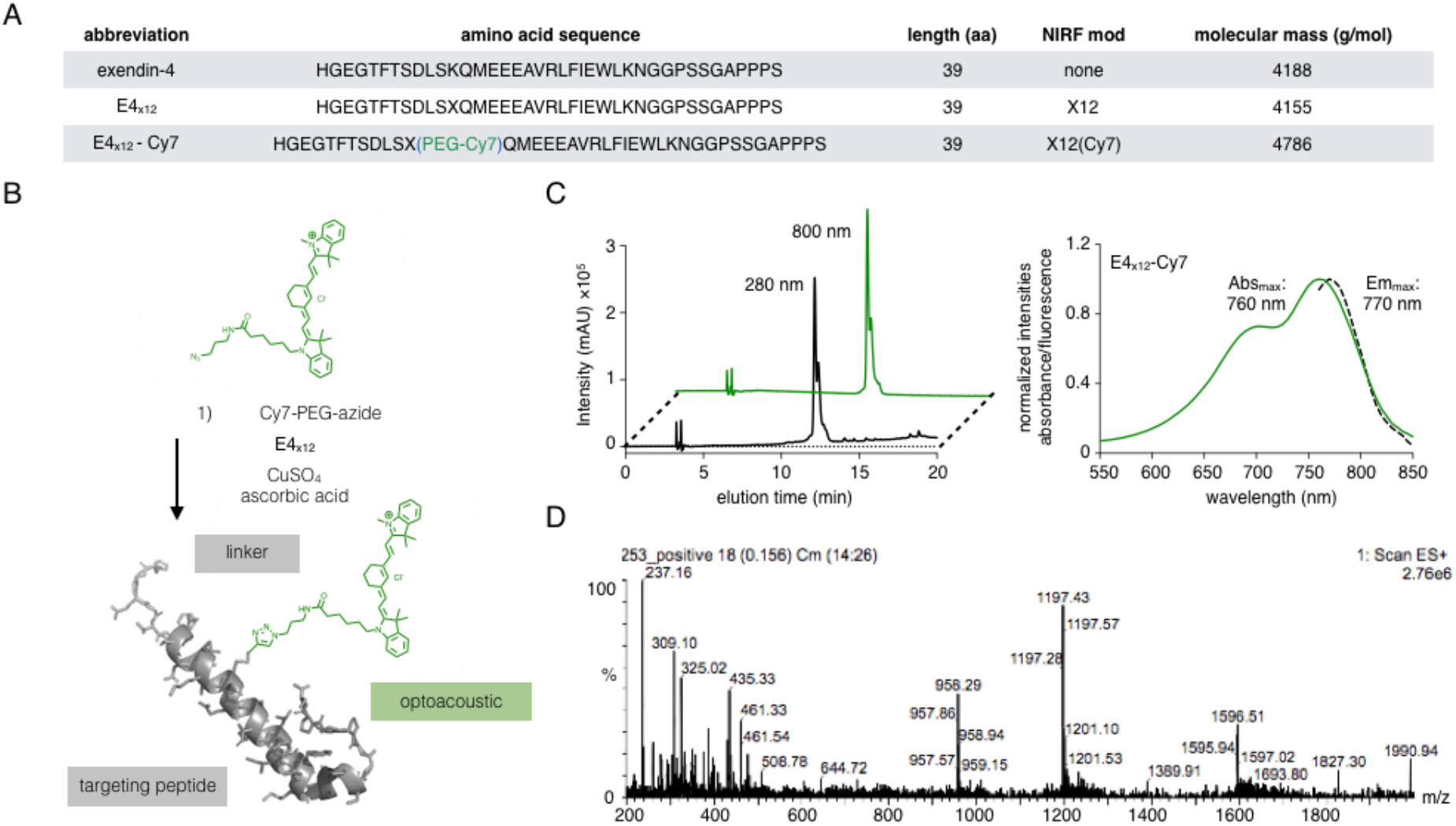
Synthesis and analysis of E4_x12_-Cy7. (A) Abbreviations and amino acid sequences of exendin-4 and its modified derivatives. (B) Copper-catalyzed azide-alkyne cycloaddition reaction scheme yielding optoacoustic probe E4_x12_-Cy7. (C) HPLC trace, 5-95% MeCN, 15 min linear gradient (*left*) and absorbance/emission spectra (*right*) of E4_x12_-Cy7. (D) ESI-MS spectrum of E4_x12_-Cy7.

### Validation of E4_x12_-Cy7 as an optoacoustic agent in tissue-mimicking phantom

The multi-spectral optoacoustic tomography (MSOT) device (MSOT inVision 256, iThera Medical) was used to measure optoacoustic signals. A flow device, fitted into the commercially available MSOT imaging system and containing a soft-tissue mimicking phantom, was prepared as previously described in detail^35, 36^. In order to evaluate the optoacoustic performance of E4_x12_-Cy7, solutions of E4_x12_-Cy7 in DMSO at concentrations between 2 μM and 25 μM) were prepared. Using a syringe pump, the samples were allowed to flow at roughly 1 mL/min until they reached the middle of the phantom. The observed optoacoustic spectrum of E4_x12_-Cy7 is broad at all of the measured concentrations (Figure 2A), with optoacoustic maxima at 760 nm, in agreement with the absorbance maxima at 760 nm (Figure 1C). The optoacoustic spectrum at 25 μM was used as the reference spectrum for MSOT spectral unmixing. After spectral unmixing, there was a linear correlation between the optoacoustic signal and concentration of E4_x12_-Cy7 (R^2^ = 0.99). Representative 2D optoacoustic images of the phantom with the E4_x12_-Cy7 at varying concentrations were reconstructed, and 900 nm was chosen as the representative optoacoustic background (Figure 2D). A linear regression fitting algorithm using classical least square of known spectra was used to unmix the raw optoacoustic data (a.u.), and the algorithm gives out a score. Hence the name CLS score^38–40^. In our case, three references were included and used for spectral unmixing of hemoglobin, deoxyhemoglobin and E4_x12_-Cy7.

**Figure 2.**
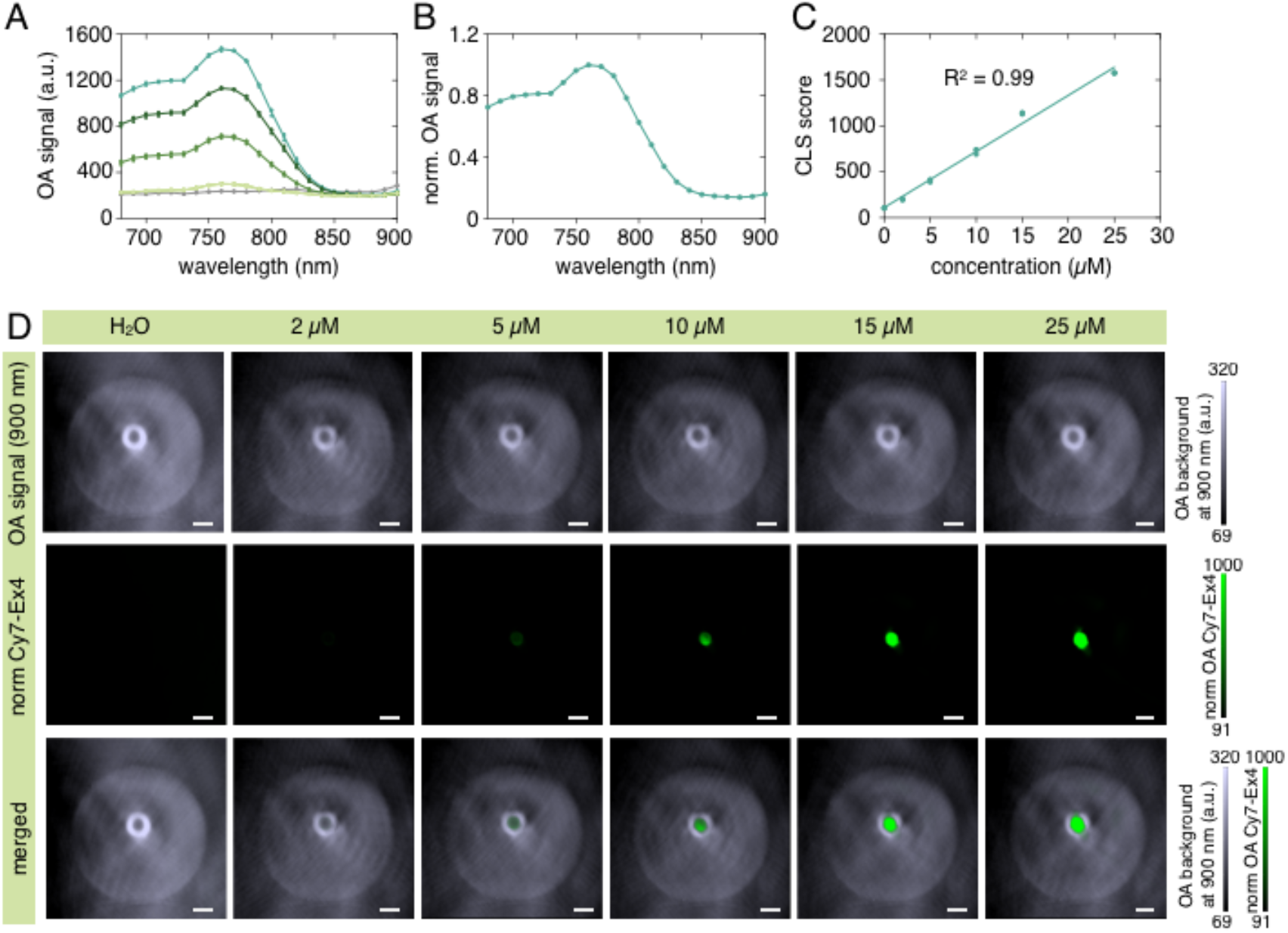
Evaluation of pancreatic ß-cell targeting peptide (E4_x12_-Cy7) as an optoacoustic probe in soft-tissue mimicking phantoms. (A) Optoacoustic spectra from 680-900 nm with 10 nm wavelength resolution at various concentrations of E4_x12_-Cy7 (2 μM, 5 μM, 10 μM, 15 μM and 25 μM). (B) Normalized spectra (25 μM) at the maxima optoacoustic signal (C) Optoacoustic signal at various concentrations obtained after multi-spectral unmixing. (D) Optoacoustic image reconstruction of E4×_12_-Cy7 solutions embedded in a homogenous and absorbing soft-tissue mimicking phantom. White scale bar is 2.5 mm.

### Binding, affinity and inhibition

Cell assays were performed using MIN6 cells and NIT-1 cells, both were derived from mouse pancreatic ß-cell insulinomas. E4_x12_-Cy7 is both an optoacoustic agent and a fluorescent agent, with an IC_50_ of 4.6 ± 0.8 nM (Figure 3A). In general, concentrations lower than 50 nM produced the best results, likely due to self-quenching of the tracer at higher concentrations. Incubation of E4_x12_-Cy7 below 50 nM concentration regimes using MIN6 cells showed a relative increase in fluorescence with increasing concentrations (Figure S1A). Incubation of E4_x12_-Cy7 at 50 nM in the MIN6 cell line prove to be the optimal concentration for fluorescence imaging (Figure 3B). To test the specificity of E4_x12_-Cy7 for GLP-1R, we co-incubated MIN6 cells non-fluorescent exendin-4 (500 nM) and varying concentrations of E4_x12_-Cy7 (Figure S1B). Co-incubation of exendin-4 (500 nM, 5-20-fold excess) relative to E4_x12_-Cy7 produced no fluorescence uptake within the cells (Figure S1B). These results show that E4_x12_-Cy7 was taken up specifically via by GLP-1R and that its uptake could be inhibited completely by the presence of excess unlabeled peptide. Binding of E4_x12_-Cy7 to the GLP-1R induced internalization resulting in cytosolic as well as membrane localization at 90 min incubation.

**Figure 3.**
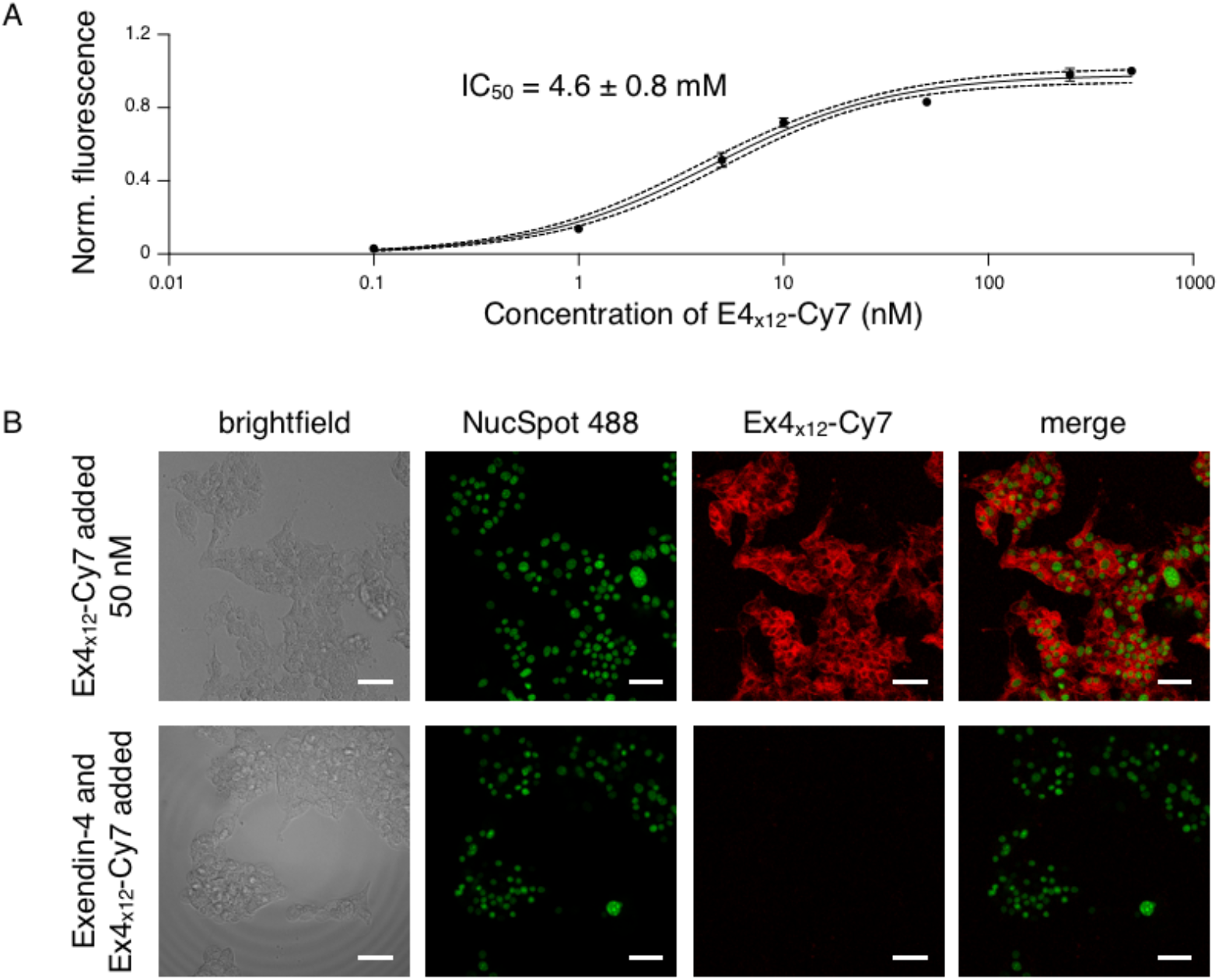
*In vitro* binding and inhibition studies. (A) IC_50_ value of E4_x12_-Cy7 measured via a competitive binding assay using NIT-1 cells. (B) Confocal microscopy imaging experiments using MIN6 cells following the addition of E4_x12_-Cy7 (top panel) or after coincubation with excess exendin-4 (lower panel). From *left* to *right* is the brightfield, nuclear staining with NucSpot 488, 50 nM E4_x12_-Cy7 staining or coincubation with excess exendin-4 and its composite. Scale bars are 40 μm.

### GLP-1R mediated accumulation of E4_x12_-Cy7

After MSOT pre-imaging sessions, mice used for blocking experiments were injected with 151 μg of exendin-4 (100 μL, 360 μM, 30 min time interval) before i.v. injection with 57 μg E4_x12_-Cy7 (100 μL, 120 μM, 30 min time interval) and MSOT imaging were carried out. Likewise, mice that were injected with saline or 57 μg of E4_x12_-Cy7 were imaged before and after 20 min injections. Ex vivo biodistribution via fluorescence imaging showing that the kidney has the highest accumulation of E4_x12_-Cy7 (Figure S3). A region of interest (ROI) drawn around the kidney before and after MSOT imaging (Figure 4A) shows that the optoacoustic spectral profile matches that of E4_x12_-Cy7 (Figure 2A/B). As a reference point, a ROI around an area that is not kidney were also plotted as background. Therefore, there can be no doubt that the signal after spectral-unmixing is from the Cy. To confirm further, the organs from the pancreas, muscle and insulinoma MIN6 were immediately harvested and imaged with the MSOT (Figure 4B). A statistically significant differences between muscle and the pancreas (p**** < 0.0001) and between the muscle and MIN6 (p**** < 0.0001) were observed (unpaired t-test). In addition, in vivo epifluorescence imaging showed statistically significant differences between the blocking and E4_x12_-Cy7 groups (p** = 0.0042), showing that the signal was blockable (Figure 4C). Moreover, organs (insulinoma MIN6, pancreas, muscle, kidney, spleen and liver) were excised and placed side-by-side on a black non-fluorescent paper. Epifluorescence imaging were carried out, an ROI were drawn around the edges of the organs, and the average fluorescence signal was calculated. The signal originating from the mouse insulinoma MIN6 is statistically significant (p* = 0.0199) between the blocking and E4_x12_-Cy7 groups (Figure 4D).

**Figure 4.**
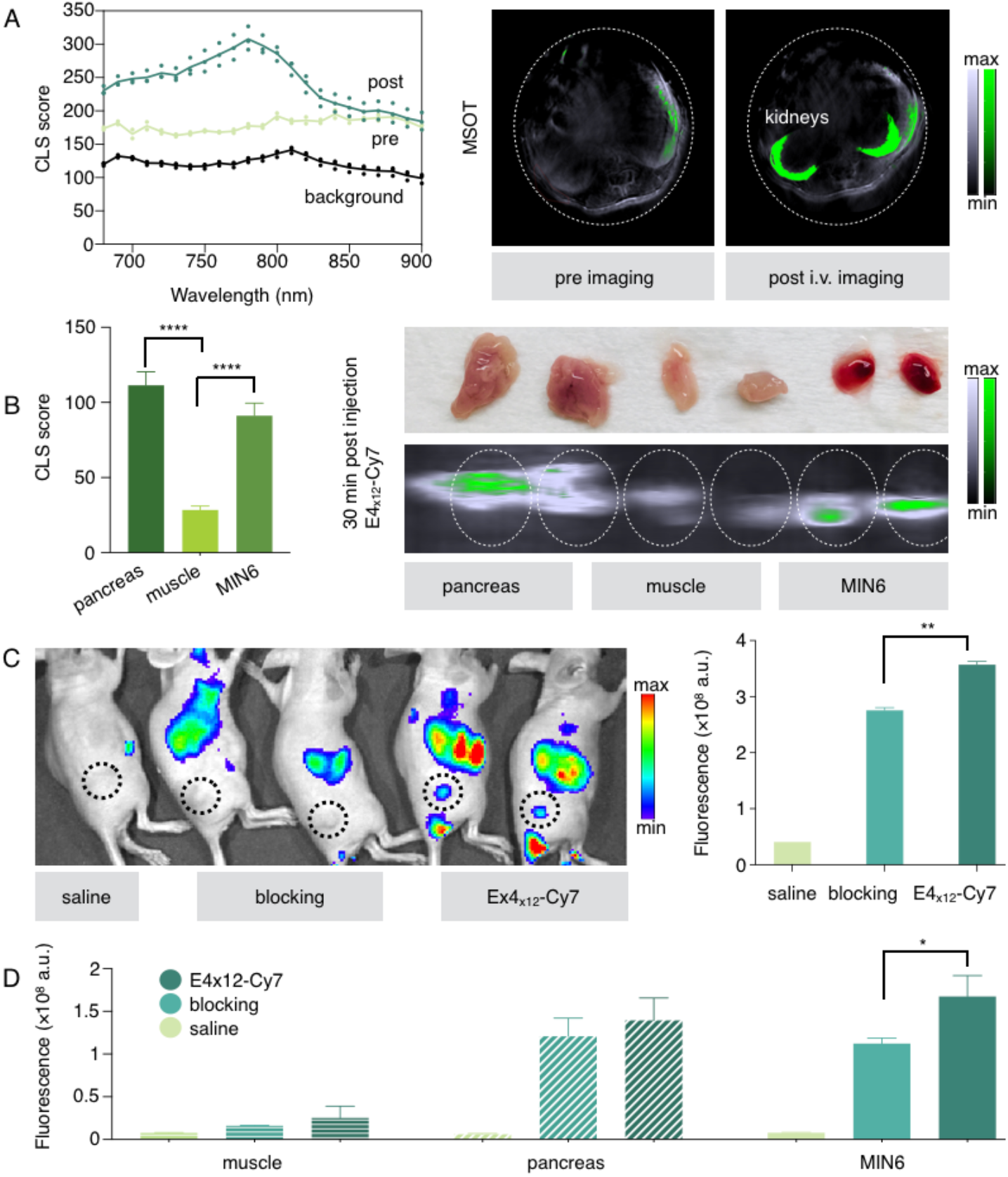
The accumulation of E4_x12_-Cy7 is GLP-1R mediated. (A) Optoacoustic signal at varying wavelengths (680-900 nm, 10 nm step) taken at the kidney region (*left*) and the corresponding MSOT images (*right*). (B) Ex vivo MSOT images of organs from mice that were injected with E4_x12_-Cy7. From left to right are the pancreas, muscle and MIN6 (*right*) and the corresponding quantification (*left*). GLP-1R mediated accumulation. (C) Representative in vivo epifluorescent images of mice that were injected either with saline, blocking or E4_x12_-Cy7 (*left*) and quantification (*right*). (D) Corresponding ex vivo epifluorescent images from left to right of MIN6, muscle and pancreas that were either injected with saline, blocking or E4_x12_-Cy7 (*left*) and quantification (*right*).

### Optoacoustic imaging of insulinoma

MIN6 cells were used in a mouse model of pancreatic insulinoma since they closely resemble pancreatic ß-cells, including their GLP-1R expression, and insulin secretion^37, 41^. To determine specific optoacoustic detection and optoacoustic imaging of insulinoma xenografts *in vivo*, mice were intravenously injected with 57 μg E4_x12_-Cy7. MSOT imaging of the same mice before injection served as the negative control. Optoacoustic imaging using the MSOT before and 30 min after administration showed a statistically significant increase in signal within the tumors (Figure 5A). Mice that were imaged 30 mins after intravenous injection of E4_x12_-Cy7 had an optoacoustic signal score of 72.69 ± 2.72 (n *= 3* mice). Pre-injected mice had an optoacoustic background signal score of 30.54 ± 1.80 (n *= 3* mice). Optoacoustic signal showed statistically significant differences in mouse insulinoma signal before and after intravenous injection of E4_x12_-Cy7 (*p = 0.0236). The signal strength ratio between the injected and non-injected mice was found to be 2.38 (Figure 5B). In addition, fluorescence imaging *in vivo* and *ex vivo* was carried out (Figure S3). Fluorescence imaging corroborated our optoacoustic imaging observations. *Ex vivo* analysis of E4_x12_-Cy7 showed uptake in the kidney and liver as well (Figure S3).

**Figure 5.**
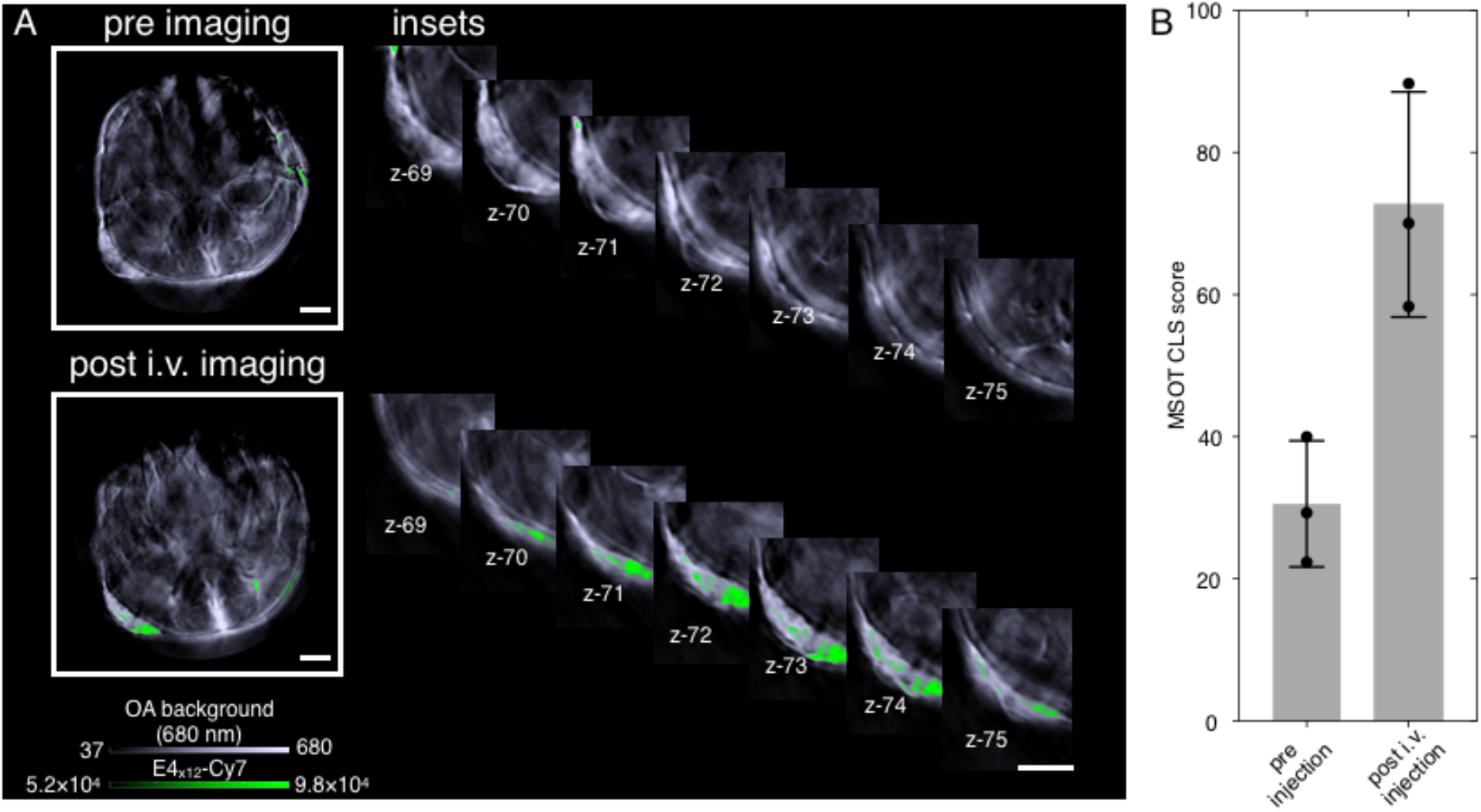
*In vivo* multi-spectral optoacoustic evaluation of E4_x12_-Cy7. (A) Optoacoustic image reconstruction showing transverse 2D projection of mice at the region of interest before and 30 min after intravenous injection of E4_x12_-Cy7 (2.9 mg/kg). Overall optoacoustic signal at 680 nm (*grey scale*) is overlaid on top of multi-spectrally unmixed signals showing the E4_x12_-Cy7 channel (*green channel*). Both scale bars are 2.5 mm. (B) Optoacoustic signal quantification after multi-spectral unmixing.

## Discussion

Optoacoustic technologies are quickly gaining increasing relevance in diagnostics, tracking disease progression, as well as evaluating treatment therapies through imaging of disease-relevant biological processes^42^. For this application, the most attractive feature of MSOT over other optoacoustic systems is its ability to provide functional information, including a range of wavelength measurements in the NIR region that enable multi-spectral unmixing between endogenous and exogenous contrast agents. Hence, a highly absorbing NIR agent, such as our E4_x12_-Cy7, can achieve sufficient contrast for *in vivo* ß-cell detection. Compared to traditional fluorescence imaging systems, the key difference of MSOT is that its ability to use light for excitation and sound for detection enables a greater penetration depth. The development of optoacoustic imaging tools for diabetes is a valuable but unmet clinical need^7, 8, 25, 31, 43^. A handful of proof-of-concept studies in the literature have emerged to show the potential of diabetes monitoring via optoacoustic imaging, but none feature direct visualizations of ß-cells^6–9, 19, 44–46^.

Recent studies typically demonstrated an indirect biological response to the symptoms of diabetes via vasculature tracking^44, 47^, diabetic neuropathy^48^ or glucose sensing^8, 45, 46^. And while CT, MRI, ultrasonography and intravital microscopy^4^ are well established in the field of preclinical diabetes imaging, the techniques are hampered by photon absorption and scattering in tissues. Here, an optoacoustic agent E4_x12_-Cy7 was successfully synthesized, purified, chemically and optoacoustically characterized and validated *in vitro* and *in vivo*. Our rationale behind Cy7 as the absorber of choice for conjugating to neopeptide E4_x12_ was its high extinction coefficient and broad absorption profile^36^, ideal for optoacoustic detection. The introduction of an optoacoustic tracer, Cy7, to the 12^th^ position of the amino acid sequence did not perturb its binding affinity to GLP-1R, similar to what we found earlier when substituting at the 14^th^ position^49^. E4_x12_-Cy7 maintained its high receptor binding and cellular internalization at 90 min post-incubation and proved to be receptor specific, since it was blocked by 5-20 fold excess exendin-4.

For *in vivo* imaging, we chose GLP-1R-expressing mouse insulinoma MIN6, derived from transgenic mice and preserving normal characteristics and function of the endocrine ß-cells^37^. *In vivo*, healthy 6-8-week-old Fox1^n^ nude MIN6-bearing mice were injected with E4_x12_-Cy7. Intravenous injection of E4_x12_-Cy7 showed specific uptake after 30 mins, which we were able to detect with both MSOT and fluorescence imaging in the same animal. Cy7 has a broad optoacoustic spectra with maxima in the near infrared (760 nm). A similar optoacoustic spectral profile was obtained in the kidney regions after i.v. injection, confirming the presence of E4_x12_-Cy7 in vivo. It was further corroborated with ex vivo MSOT imaging of the corresponding organs of interests. This signal was blockable with exendin-4. To test E4_x12_-Cy7, *in vivo* imaging was performed in mice with subcutaneous MIN6 insulinomas. This model was chosen to focus on reproducibility, reducing the complexity that comes with imaging the pancreas. Initially, the visualization of a solid subcutaneous ß-cells is much easier than the visualizations of the 1-2% ß-cells that are spread throughout the pancreas. Moreover, we are minimizing the physiologic challenges that exists when imaging mouse pancreata. A mouse pancreas is rather flat and dendritic-like while in humans, it is a well-defined solid mass. Nevertheless, we have also demonstrated uptake in the pancreas. From our model, we have observed rapid clearance after 30 min post i.v. injection of E4_x12_-Cy7, possibly due to the short half-life (1-2 minutes) of GLP-1 caused by N-terminal degradation by the enzyme dipeptidyl peptidase-4 (DPP-4). Also, the high extinction coefficient of Cy7 dye (206, 000 M^−1^ cm^−1^) may not be adequate and we suspect its fluorescent nature may not be optimal as an imaging agent for MSOT. Taken together, this could potentially be a limitation and leaves room for further development of other optoacoustic ß-bell targeting agents.

Exendin-4 allows for modular design without perturbing its affinity to GLP-1R. Here, we have shown the first development and *in vivo* validation of a NIR MSOT agent, E4_x12_-Cy7, an exedin-4 analog that binds to GLP-1R. The MSOT can achieve multi-spectral unmixing methods which delivers information specific from E4_x12_-Cy7. Consequently, E4_x12_-Cy7 could work well with the hand-held optoacoustic imaging system. In addition, the incorporation of 3D optoacoustic tomographic systems could work in favor with near infrared absorbers, known to provide more quantitative assessments as it has greater sensitivity than 2D systems^30^. Based on the previous success of specific ß-cell imaging agents for PET, fluorescence imaging or in combinations thereof ^4, 24, 26^, we envision that a contrast agent such as E4_x12_-Cy7 could represent a potential platform for further development and optimization of ß-cell optoacoustic imaging.

## Conclusion

In this study, we synthesized and characterized E4_x12_-Cy7, an optoacoustic imaging agent based on exendin-4, a first generation GLP-1R-targeted optoacoustic ß-cell agent. Using MSOT, E4_x12_-Cy7 showed specific uptake for MIN6 xenografts. The high spatial resolution and penetration depth achieved with MSOT, in combination with targeted NIR E4_x12_-Cy7, could represent a valuable technique for monitoring diabetes and its progression in both preclinical and clinical contexts. This study is the first step towards the development of ß-cell mass optoacoustic imaging, a technique that merits further investigation as the already enormous challenge diabetes represents continues grow.

## Materials and Methods

### General

Liquid chromatography-mass spectrometry (LC-MS) using electrospray ionization (ESI) was recorded using a Waters instrument with SQD detector for mass identification. A lyophilizer (FreeZone 2.5 Plus, Labconco, Kansas City, MO, USA) was used for freeze drying. A BS-8000 120 V high pressure (Braintree Scientific, Inc., MA, USA) multi-syringe pump was used to control the flow rate and facilitate delivery of solutions inside the phantom flow system. An automated cell counter (Beckman Coulter, Vi-Cell viability analyzer) was used for counting the number of cells. *In vitro* fluorescence confocal microscopy was carried out using a Leica TCS SP8. *In vivo* and *ex vivo* fluorescence imaging was carried out using a planar (2-D) IVIS Spectrum device (PerkinElmer, Waltham, MA, USA). Pymol (1.7) was used for the 3D peptide visualization. Exendin-4 structure (ID #3C59) was taken from RCSB protein data bank (PDB).

### Chemicals

All materials were obtained from Sigma Aldrich (Milwaukee, WI), unless otherwise specified. All reagents were used without further purification. Wild-type exendin-4 (H-HGEGTFTSDLSKQMEEEAVRLFIEWLKNGGPSSGAPPPS-NH_2_) was purchased as a custom peptide from Innopep (San Diego, CA). Single mutant exendin-4 (E4_x12_) (H-HGEGTFTS**X** LSKQMEEEAVRLFIEWLKNGGPSSGAPPPS-NH_2_), where **X** is the non-natural amino acid (S)-2-amino-4-pentynoic acid, was custom synthesized from CSBio (Menlo Park, CA). Cy7 azide (no sulfate groups) was purchased from Lumiprobe (Hallandale Beach, FL).

### Preparation of E4_x12_-Cy7

Preparation and characterization of Cy7 single mutant exendin (hereon referred to as E4_x12_-Cy7) was performed using azide-alkyne click chemistry. Briefly, 300 nmol of Cy7 azide was added to 100 μL of 1:1 water/*tert*-butanol containing 3 μL CuSO_4_-THBTA (100mM in water) and 15 μL L-sodium ascorbate (50 mM in water). Lastly, single mutant exendin E4_x12_ was added in an equal molar ratio (300 nmol) as the Cy7 azide. The mixture was reacted under magnetic stirring for 3 hours at room temperature (20–25 °C). Purification was performed on a Shimadzu RP-HPLC system equipped with a DGU-20A 5R degasser, SPD-20AV UV/Vis detector, a LC-6AD pump system, CTO-20A oven maintained at 30 °C, and a CBM-20A communication BUS module and using a C-18 solid phase analytical column with mobile phase of water (0.1% trifluoroacetic acid, Buffer A) and acetonitrile (0.1% trifluoroacetic acid, Buffer B). A 30– 80% CH_3_CN method (gradient: 0–18 min 30–80%, 18-20 min 80-30%) at a flow rate of 1 mL min^−1^ was used to separate E4_12_-Cy7 (~16min) from free Cy7 azide (~21 min) and unlabeled E4_x12_ (~12 min). Purified samples were lyophilized overnight, reconstituted in 1:1 water/acetonitrile and stored at −20 °C until further use. Molecular weight of the purified product was confirmed using electrospray ionization (ESI) (calculated 4786.39; found 4785.36).

### Cell culture

Mouse insulinoma-derived MIN6 were used between passage 13 and 17. Cells were grown on in high-glucose DMEM containing 15% vol/vol heat-activated FBS, 50 U/mL penicillin, and 10 μg/mL streptomycin. GLP-1R positive NIT-1 mouse cells were cultured in F12K media supplemented with 10% (v/v) FBS, 50 U/mL penicillin, 50 μg/mL streptomycin, and 1.5g/L sodium bicarbonate.

### *In vitro* binding affinity assay

Confluent NIT-1 cells were harvested using 0.05% trypsin-EDTA for 5 minutes before the enzymes were neutralized by adding FBS containing F12K media. Cells were centrifuged gently at 300 rpm for 5 min at 4 °C, and re-suspended in 2 mL PBS containing 0.5% bovine serum albumin (BSA). Cells were aliquoted and incubated in triplicate with 0.5% BSA/PBS containing E4_x12_-Cy7 at varying concentrations (0.1 nM – 250 nM) for 3 hours on ice. A control for non-specific bindingwas included by pre-incubating cells with 2 μM of unlabeled wild-type exendin on ice, followed by 0.5% BSA/PBS containing 250 nM E4_x12_-Cy7 and 2 μM for 3 hours on ice. Cells were washed with 0.5% BSA/PBS and analyzed by flow cytometry (Bio-Rad ZE5 Cell Analyzer). Binding affinity (IC_50_) was estimated using GraphPad Prism 8 using a one-site specific binding model.

### Cell imaging

MIN6 (1×10^5^ cells/well) cells were seeded into a 96-well plate and cultured for 72 h (37 °C). The cells were then incubated with various concentrations of E4_x12_-Cy7 for 90 min at 37 °C alone or with exendin-4 (exenatide; 500 nM) co-incubated with various concentrations of E4_x12_-Cy7 (25 to 100 nM) for 90 min at 37 °C. Following incubation, the cells were washed once with 200 μL PBS and fixed by adding 100 μL of 4% PFA and allowed to sit at room temperature for 15 min. PFA was removed and washed with PBS (200 μL, 3×). The cells were mounted on PBS with NucSpot 488 and the cells were imaged with confocal fluorescence microscope (Leica TCS SP8).

### Tissue mimicking phantom preparation and experimental setup

A phantom mold was prepared to fit the MSOT holder. The phantom allows for a flow-mediated setup as previously shown^35^. Briefly, a clear, cylindrical hollow membrane tubing was placed at the center of a cylindrical mold (20 mL syringe, diameter: 2 cm, length: 14 cm). The soft-tissue mimicking phantom was freshly prepared by adding 15% v/v intralipid® 20%, I.V. fat emulsion (to provide the scattering), and 0.01 mM Direct Red 81 (for absorption) to a pre-warmed solution of 5% v/v agarose Type 1 (solid in <37 °C) in Milli Q water (18.2 MΩ cm^−1^ at 25 °C). At 15% agarose, the phantom was suspended in water with the weight of the hollow tubing being supported. The mixture was poured into the mold and allowed to cool and solidify for at least 2 hours and up to overnight in the fridge. At each end of the tubing, a luer-lock was fitted to connect the syringe pump, equipped with a 20 mL syringe filled with PBS. Air gaps were used to separate the sample and PBS. The flow rate was kept at 1 mL min^−1^ until the sample of interest reached the region of interest (ROI) for imaging. After imaging, the flow rate was increased 2× and the phantom was washed with at least 10 mL PBS. The wash, load sample and wash cycle were repeated for all samples. The flow-mediated phantom setup sustained several acquisitions throughout the day and the same phantom setup was used for all samples that were compared to each other.

### Multi-spectral optoacoustic tomography (MSOT)

A multi-spectral optoacoustic tomography (MSOT) inVision device (MSOT inVision 256, iThera Medical, Munich, Germany) for small animal imaging was used. MSOT inVision has an array of 256 detector elements that are cylindrically focused, with a central ultrasound frequency of 5 MHz and up to 270° coverage. The soft-tissue mimicking phantoms were aligned so that the illumination ring coincided with the detection plane, i.e. the curved transducer array was centered around the phantom. Data were obtained in the wavelength range of 680–900 nm (10 nm increments) and an average of 10 frames per wavelength were acquired, which equates to 1 s acquisition time per wavelength per section. The optical excitation originates from a Q-switched Nd:YAG laser with a pulse duration of 10 ns and a frequency of 10 Hz. Light is homogenously delivered to the phantom using a fiber split into 10 output arms. The fiber bundle and the transducer array are stationary, and the sample holder moves along the z-direction allowing longitudinal acquisition of different imaging planes using a moving stage. MSOT measurements were performed in a temperature-controlled water bath at 34 °C. All variable parameters such as optoacoustic gain, laser power, focus depth, frame averaging, frame rate and high/low pass filters were kept constant during the measurement. Samples were allowed to equilibrate for a minimum of 5 min before initiating the optoacoustic scan, and concentrations of E4_x12_-Cy7 ranged from 2 μM, 5 μM, 10 μM, 15 μM and 25 μM in 0.9% sodium chloride. Sample solutions were delivered to the center of the phantom using a syringe pump. Between the measurements, the flow device was washed with deionized water (> 10 mL).

### Optoacoustic image data processing

Spatial reconstruction and multi-spectral processing (MSP) of the data was performed using the ViewMSOT software suite (V3.6; iThera Medical) and a backprojection algorithm. The normalized optoacoustic spectra in Figure 2B were obtained from optoacoustic phantom scans taken from a 25 μM sample and used as a contrast agent reference. For multi-spectral unmixing, a linear regression method was used and excluded pixels where the number of wavelengths (> 25%) fell within the negative signal. Hemoglobin (HbO_2_) and deoxyhemoglobin (Hb) reference spectra included in the software package were used for multi-spectral unmixing. MIN6 cells were implanted in the same position and distance from the surface of the skin across animals. Hence, all animals are affected to the same degree by light scattering. Quantitative information was obtained by defining the ROI manually within a 2D MSOT image. For spectrally unmixed E4_x12_-Cy7, the mean pixel intensities per cross-section in the volume of interest (VOI) were first plotted as classical least square (CLS) score vs. position (mm) to assess its signal strength. Signal strength decreasing with distance from the xenograft and creating a parabolic shape agress with the analysis. The maximum “mean signal per cross-section” was used as a quantitative indicator of the probe binding.

Overall optoacoustic intensities (a.u.) were reported at all imaging wavelengths (680– 900 nm, 10 nm step resolution) (Figure 2A). For all cases, unless otherwise stated, the (CLS) method for spectral deconvolution was used and reported as CLS scores. The following reference spectra were used: hemoglobin (HbO_2_), deoxyhemoglobin (Hb) and E4x12-Cy7. To compare the direct intensities of all images, the optoacoustic spectra (25 μM) were normalized to their maximum optoacoustic (760 nm) wavelength. The channel for E4_x12_-Cy7 (*green*) after multi-spectral unmixing was overlaid with the overall optoacoustic intensities (a.u., *bone scale*).

### Animal studies

All animal experiments were performed in accordance with protocols approved by the Institutional Animal Care and Use Committee (IACUC) of Memorial Sloan Kettering Cancer Center (MSK) and followed the National Institutes of Health guidelines for animal welfare. Healthy Hsd:athymic female mice NudeFoxn1^nu^ (6–8 weeks old) were obtained from the Jackson Laboratory. The mice were split into two groups of three mice, one group injected with the imaging agent and the other used a non-injected control group. All animal procedures, other than tail vein injections, were performed with the animals under general 1.5-2% isoflurane inhalation anesthesia. In order to test GLP-1R receptor specificity, xenografts were created using the pancreatic mouse insulinoma line MIN6. MIN6 cells were injected into the right flank (1 × 10^6^ cells in 50 μL 1:1 HG DEM medium and Matrigel). The xenografts were left for 12–15 days to allow for proper vasculature formation around the ß-cell clusters prior to imaging. From MSOT imaging, the mice were placed into the animal holder in supine position, gently fixed into position using clear straps and fitted with a breathing mask for anesthesia. Ultrasound colorless gel (approximately 1–2 mm thick layer) was applied onto the mouse around the region of interest in order to improve acoustic coupling. The animal holder was closed, wrapping the clear plastic membrane around the mouse and air gaps and bubbles in between the membrane and mouse's skin were removed. The animal holder with the mouse positioned inside it was then placed into the imaging chamber, with the animal being aligned with regards to the detection plane (centered within the curved transducer array). MIN6-bearing mice (n = 3) were imaged with the MSOT prior to injection. Group 1 mice was injected via tail vein with 100 μL of 2.9 mM kg^−1^ of E4_x12_-Cy7. The mice were re-imaged at 30 min post injection. E4_x12_-Cy7 was dissolved in 0.9% sodium chloride solution. Animals were also imaged in tandem using an IVIS Spectrum (PerkinElmer, Waltham, MA, USA) system for planar fluorescence imaging.

### Blocking in vivo experiments

MIN6-bearing mice were split into three groups namely, saline, blocking and E4_x12_-Cy7 injected groups (n = 3). Saline group were injected with 100 μL of 0.9% sodium chloride, blocking group was injected with 151 μg of exendin-4 (100 μL of 360 μM) and allowed to circulate for 30 mins, followed by the injection of 57 μg E4_x12_-Cy7 (100 μL of 120 μM) and E4_x12_-Cy7 group was injected with 100 μg of E4_x12_-Cy7 (100 μL of 120 μM). Mice were imaged before and after injections using epifluorescence IVIS Spectrum system and the MSOT.

### Ex vivo imaging of tissues

After 30 min post-intravenous injection and optoacoustic imaging, kidney, liver, pancreas, spleen, muscle and MIN6 cluster were harvested and planar fluorescence imaging was carried out.

### Statistical analysis

Paired t-tests were performed to determine statistical difference *in vivo* before and after injection. Levels of significance are as follows: ns = not significant, *p < 0.05, **p < 0.01, ***p < 0.001 and ****p < 0.0001. Data is presented as mean ± SD.

## Supporting information

2020_Roberts_SI_optoacousticImagingGLP1R

## Acknowledgements

The authors thank the support of Memorial Sloan Kettering Cancer Center's Animal Imaging Core Facility, Radiochemistry & Molecular Imaging Probes Core Facility, Molecular Cytology Core Facility, Nuclear Magnetic Resonance Analytical Core Facility and the University of Michigan Flow Cytometry Core Facility. The authors thank Christopher Wittman for assistance in cell culture during the revision of the paper and Garon Scott for proof-reading the manuscript. This work was supported by National Institutes of Health grants R01 CA204441 (T. R.), R01 CA183953 (J.G.), R35 GM128819 (G.M.T.) and P30 CA008748.

## Author Contributions

S.R. and T.R. designed the study. S.R. and E.K and T. N. performed experiments. C.C. maintained cell culture work. S.R. and T.R. primarily analyzed, interpreted and wrote the manuscript. J.G. and G. M. T. consulted and edited the manuscript. All authors have read, provided feedback on, and approved the manuscript for publication.

