## Supplementary material for "Optoacoustic imaging of GLP-1 Receptor with a near-infrared exendin-4 analog": 2020_Roberts_SI_optoacousticImagingGLP1R

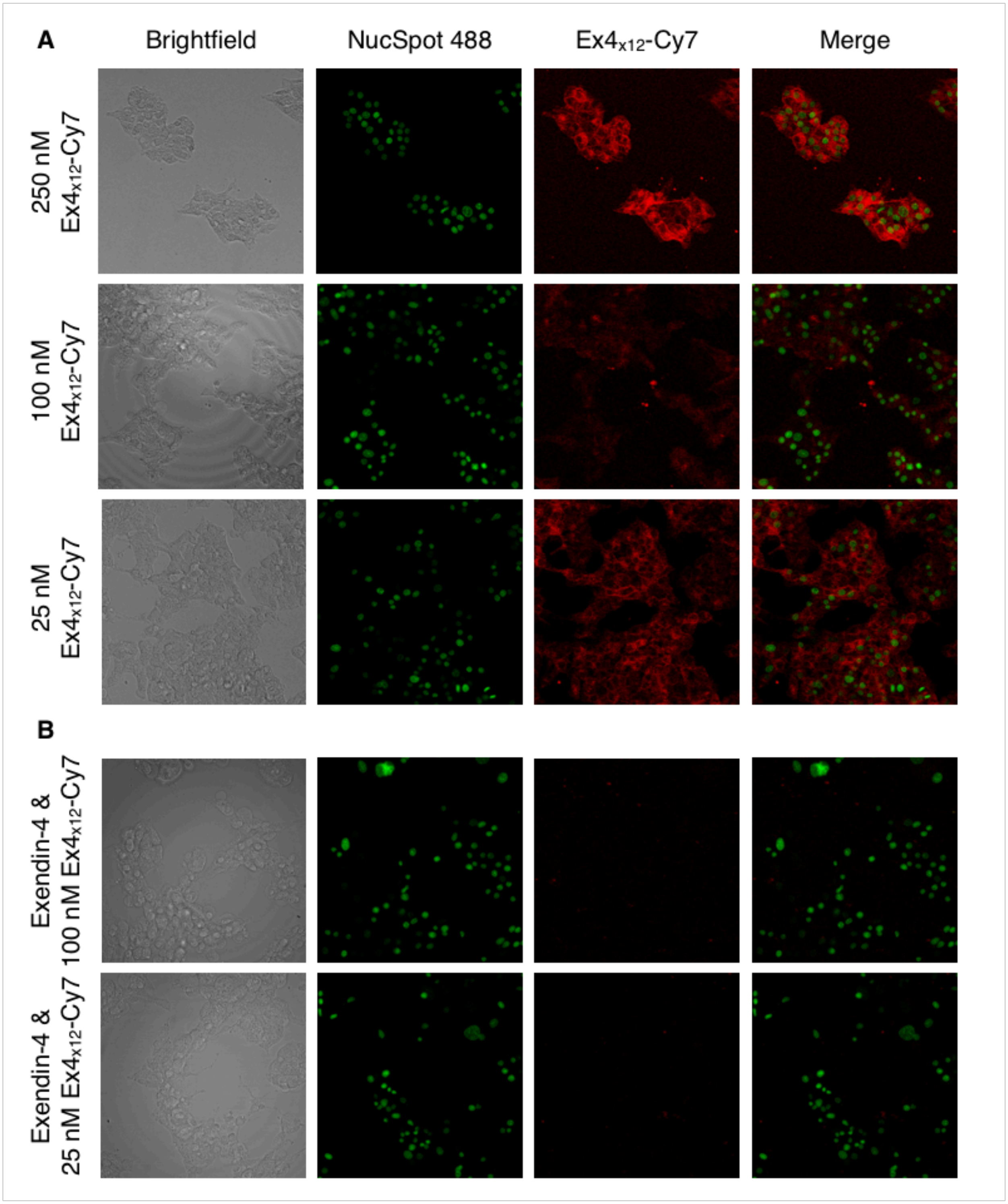


**Figure S1**. *In vitro* binding and inhibition studies. (A) Confocal microscopy imaging experiments using MIN6 cells following the addition of $E4_{x12}$-Cy7 (red) at different concentrations: 250 nM (*top row*), 100 nM (*middle row*) and 25 nM (*bottom row*).

Columns from *left* to *right:* brightfield (1st column), nuclear staining with NucSpot 488 (green, 2nd column), $E4_{x12}$-Cy7 staining (red, 3rd column) and the corresponding composite image (4th column). (B) Coincubation of $E4_{x12}$-Cy7 at 100 nM (*top*) and 25 nM (*bottom*) with excess exendin-4.

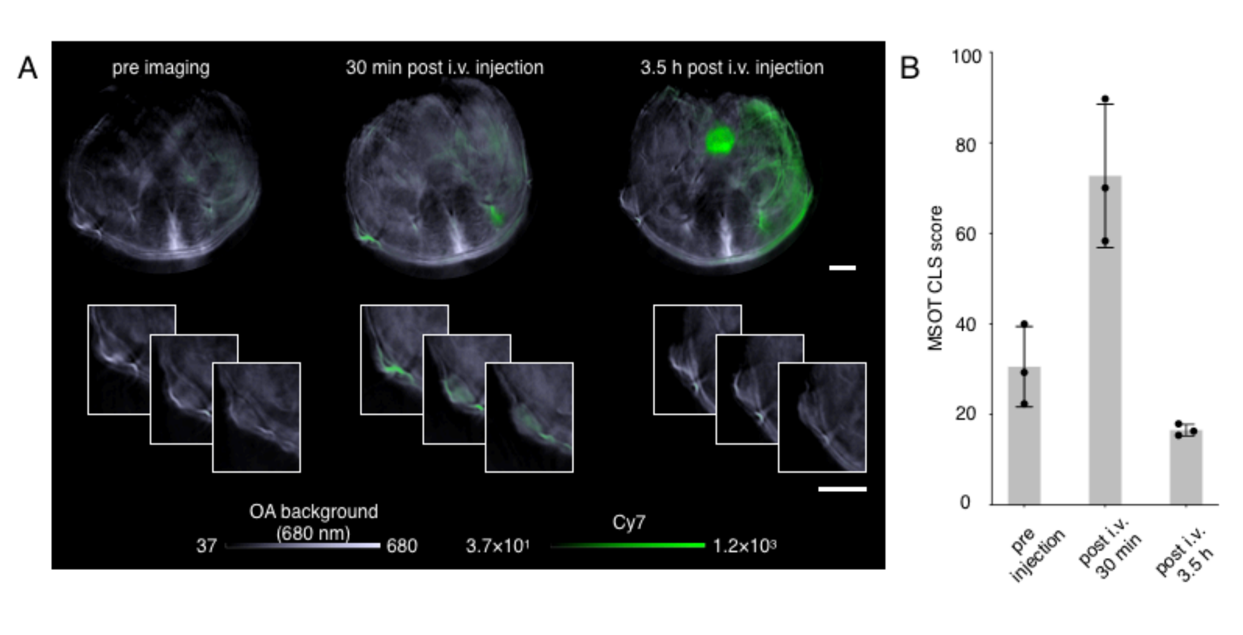


**Figure S2**. *In vivo* multi-spectral optoacoustic evaluation of $E4_{x12}$-Cy7 at different timepoints. (A) Optoacoustic image reconstruction showing transverse 2D projection of mice at the region of interest before and 30 min after intravenous injection of $E4_{x12}$-Cy7 (6.7 mg/kg) before (*left)*, 30 min post-i.v. injection (*middle*) and 3.5 h post-i.v. injection (*right*). Overall optoacoustic signal at 680 nm (*greyscale*) is overlaid on top of multi-spectrally unmixed signals showing, $E4_{x12}$-Cy7 channel (*green channel*). Both scale bars are 2.5 mm. (B) Optoacoustic signal quantification after multi-spectral unmixing.

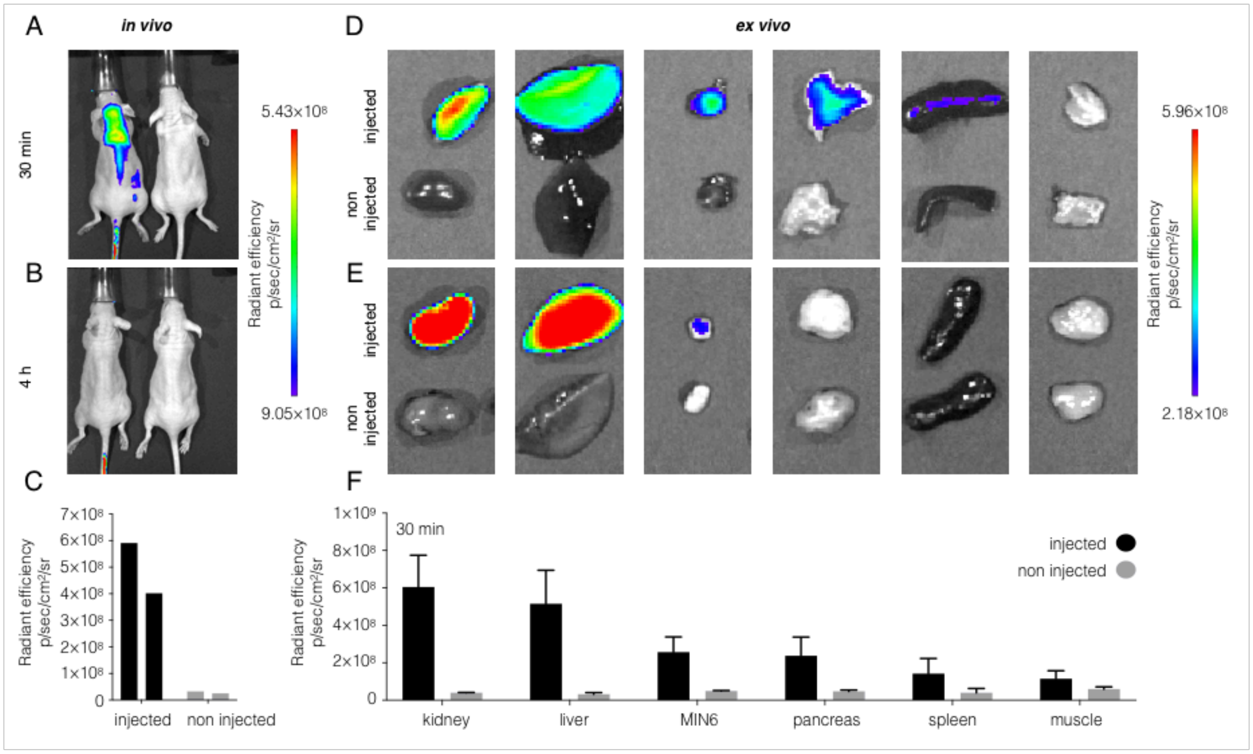


**Figure S3.** *In vivo* and *ex vivo* kinetics and biodistribution validation of $E4_{x12}$-Cy7 using fluorescence IVIS imaging. (A) Representative *in vivo* fluorescent images of $E4_{x12}$-Cy7 30 min post-i.v. injection, (B) 4 h post-i.v. injection and (C) quantifications (n = 3). (D) Corresponding *ex vivo* fluorescent images from left to right of kidney, liver, MIN6, pancreas, spleen and muscle tissues excised at 30 min and (E) 4 h post-i.v. injection of $E4_{x12}$-Cy7. (F) Quantification of the *ex vivo* organs by drawing regions of interest (ROIs) around the tissue outlines using the white field images.

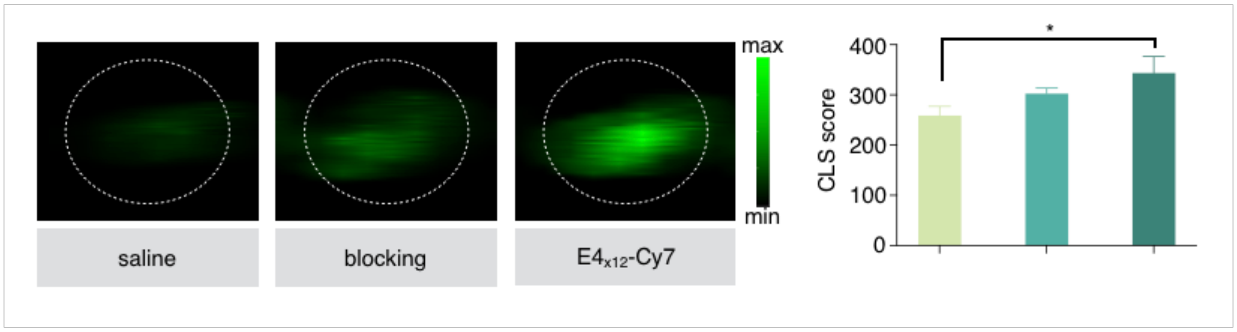


**Figure S4.** The ex vivo optoacoustic signal of kidneys between non injected, blocking and injected. From left to right are the MSOT images of the kidney from a mouse that was injected with saline, exendin-4 (151 µg) followed by the injection of $E_{4x12}$-Cy7 (57 µg, 20 mins time interval (blocking), and 57 µg of $E_{4x12}$-Cy7 (*left*) and the corresponding quantification (*right*). There is a statistically significant differences between mice that were injected with saline and $E4_{x12}$-Cy7 (p* = 0.0178).
